# CiliaBuilder2: a flexible ChimeraX bundle to visualize cilia and centriole structures in 3D

**DOI:** 10.64898/2026.10.15.759410

**Authors:** Tengqiushi Liu, Siddhartha Chakraborty, Khanh Huy Bui

## Abstract

Centrioles and cilia are large organelles essential for cell division, motility, and signaling. They are characterized by distinctive, nine-fold symmetric microtubule doublets or triplets decorated with periodic protein complexes. While cilia and centrioles are highly conserved across eukaryotes, their composition and structure vary considerably between species and tissue types. Advances in cryo-electron tomography and single-particle cryo-electron microscopy have recently led to a surge in structural data for ciliary components. However, visualizing these components within the ciliary context remains challenging due to their scale and the diverse data formats involved such as density maps, atomic models, and 3D geometric objects. Here, we introduce CiliaBuilder2, a toolset integrated into UCSF ChimeraX that enables simple, flexible visualization of complete ciliary structures from diverse types of data using parametric calculation.

## Introduction

Cilia are conserved, microtubule-based organelles that extend from the cell surface and participate in motility, signaling, and sensory functions. Motile cilia have a canonical 9+2 axoneme, in which nine outer doublet microtubules surround a central pair (Klena and Pigino 2022). The doublet microtubule has 96 nm repeat, with periodically arranged complexes, such as dynein arms, radial spokes, and the nexin–dynein regulatory complex (Bui, Yagi et al. 2012, Leung, Sun et al. 2025). Together with the central-pair apparatus that has 32-nm repeat (Oda, Yanagisawa et al. 2014, Dai, Ichikawa et al. 2020, Zhu, Lin et al. 2025), these components form an intricate structure that extends over micrometers while containing features organized on nanometer scales. Measuring about 300 to 500 nm in length, centrioles consist of nine triplet microtubules and are structurally divided into proximal, core, and distal regions (Ruehle, Li et al. 2024). Interestingly, although the cilia continue from the centrioles, the in-plane angle of the triplet microtubules and doublet microtubules are noticeably different.

Structural studies have revealed increasingly detailed views of this architecture. Cryo-electron tomography (cryo-ET) and subtomogram averaging have resolved repeating complexes in their axonemal context (Bui, Yagi et al. 2012, Oda, Yanagisawa et al. 2014, Legal, Parra et al. 2023), while single-particle cryo-electron microscopy (cryo-EM) has enabled higher-resolution reconstructions and atomic models of many components (Ma, Stoyanova et al. 2019, Kubo, Black et al. 2023, Leung, Sun et al. 2025). Research across species and mammalian cell lineages demonstrates that axoneme composition and structure differ not only among organisms and cell types, but also along a single cilium, precluding the use of a universal structural model (Bui, Sakakibara et al. 2009, Bui, Yagi et al. 2012, Leung, Sun et al. 2025). Expansion microscopy and other fluorescence imaging approaches add localization information for components that may not yet have complete structural models (Laporte, Gambarotto et al. 2024) .

These data are difficult to assemble into an organelle-scale view. Published structures are commonly distributed as separate density maps, atomic coordinate models, or other three-dimensional objects. They may describe different species, cell types, sample preparations, or limited regions of an axoneme. Integrating them requires decisions about coordinate systems, scale, orientation, component identity, and longitudinal repeat phase. Missing components must also be represented without implying that their positions or structures have been experimentally determined. Consequently, constructing a coherent ciliary scene from available data can involve extensive manual placement and revision.

Large-scale biological visualization tools demonstrate the value of combining structural information in a common scene. cellPACK generates mesoscale models from biological and structural information, while cellVIEW supports interactive visualization of large molecular assemblies (Johnson, Autin et al. 2015, Le Muzic, Autin et al. 2015). Within the popular molecular visualization software ChimeraX (Goddard, Huang et al. 2018, Pettersen, Goddard et al. 2021), ArtiaX provides tools for viewing tomograms, handling particle lists, and constructing a model of the cellular landscape by placing the subtomogram averages into its location in the tomogram (Ermel, Arghittu et al. 2022, Roth, Ermel et al. 2025). These tools address important general modeling and cryo-ET workflows. Building a cilium from components reported across multiple studies, however, also calls for a workflow organized around ciliary geometry and its repeating architecture. Such a scene can aid teaching and scientific communication and allow researchers to inspect the geometric plausibility of proposed component arrangements.

Here, we present CiliaBuilder2, a ChimeraX bundle for the parameterized construction and visualization of ciliary and centriolar architectures. The tool generates coordinate models from adjustable geometric parameters and uses them to place repeated components, reducing the need to duplicate and position each model manually. By working within ChimeraX, it can bring density maps, atomic models, and three-dimensional objects into a shared visual scene. CiliaBuilder2 is intended to make multiscale ciliary models easier to construct, revise, and communicate while keeping the distinction between experimentally derived structures and schematic components clear.

## Results

### CiliaBuilder2 Implementation

Inspired by ArtiaX (Ermel, Arghittu et al. 2022) implementation of placing averaged maps into particle coordinates in the form of a particle list file such as RELION STAR file (Burt, Toader et al. 2024), CiliaBuilder2 uses parametric inputs to generate RELION STAR file internally and calculated parametrically for the coordinate and orientation of microtubules, central pair and also peripheral complex such as Intraflagellar transport (IFT) complex (Fig. 1A, B). Instead of requiring the users to manually create every repeated copy of a structure, the program first uses the data in the STAR file which contains parameters of particle positions, orientations, pixel size and filament names. These coordinates can be rendered as markers and arrows as placement targets for maps and models.

**Fig. 1.**
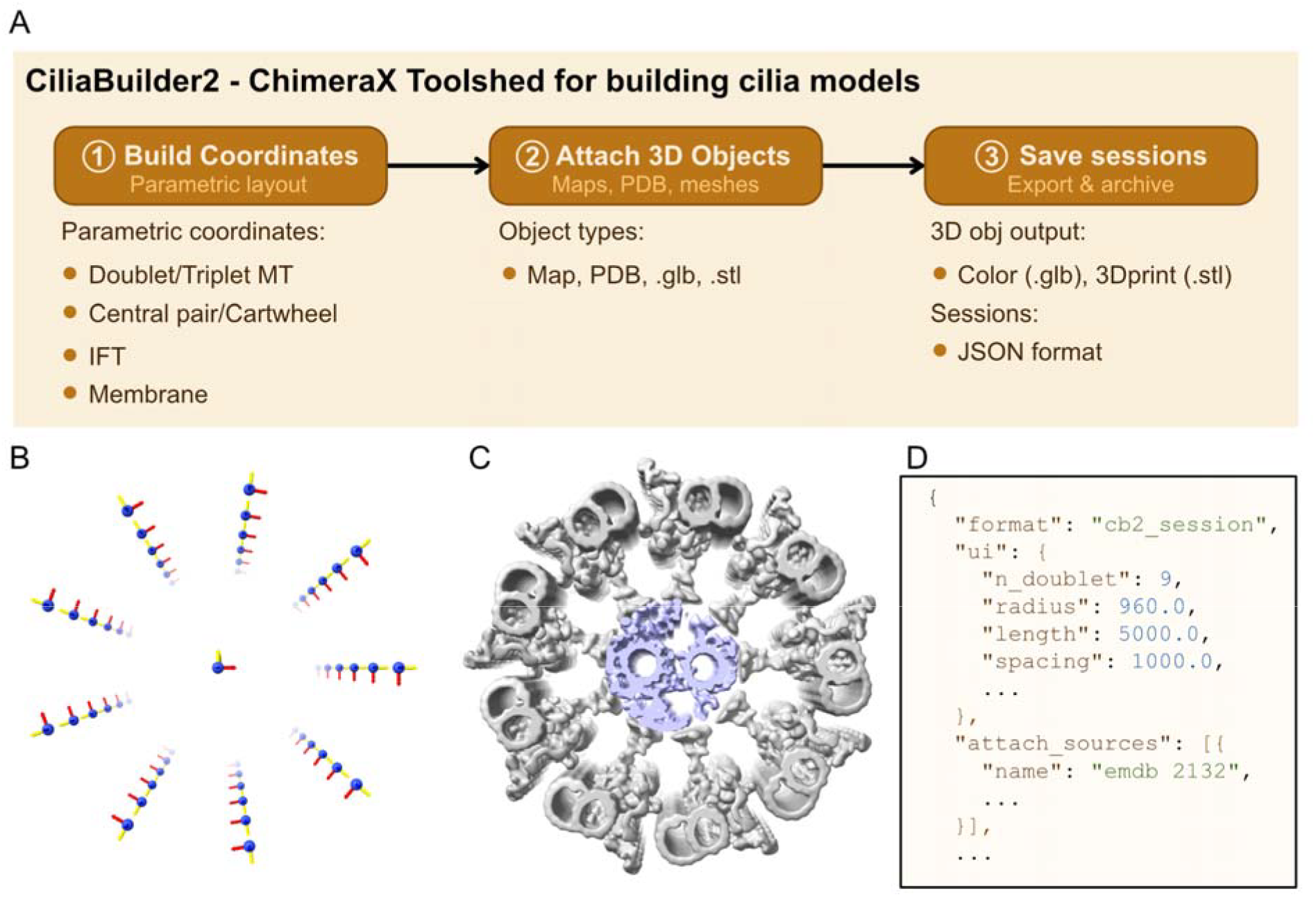
Principles of CiliaBuilder2. A. The pipeline for CiliaBuilder to build a model include (1) Build Coordinates; (2) Attach 3D Objects; and (3) Save session as JSON or 3D format. (B, C, D) Illustrations of the three steps described above.

Then users can load in a model (map, atomic model or 3D objects) and attach to the coordinate to represent multiple copies of the same object such as the 96 nm repeat of the nine doublet microtubules or the 32 nm repeat of the central pairs (Fig. 1A, C). CiliaBuilder2 allows tweaking of model or map orientation around the Y and Z axis as many subtomogram average models are generated so that the filament axis is along Y instead of Z.

In this case, our implementation is similar to the ArtiaX particle list and attaching a geometric model with flexibility of orienting the geometric model interactively. Furthermore, CiliaBuilder2 also works for PDB, GLB and STL files, which are not supported by ArtiaX. The ability to work with 3D object formats (GLB or STL) allow us to model colored maps and simple drawing of complexes that do not have structure available. As the center of rotation calculated for map is different from PDB, GLB or STL files in ChimeraX, we provided a customized center of rotation so that a cilium can be built exactly from maps or GLB or PDB that are aligned with the corresponding maps.

The complete 3D scene can be saved as a single GLB file natively supported by ChimeraX and then compressed outside to reduce file size as ChimeraX only supports reading compressed GLB but not writing compressed GLB. GLB files can be viewed on the web and imported into 3D modelling software such as Blender (Team 2022). Furthermore, we introduce a CiliaBuilder2 JSON format allowing us to save the cilium model and load again later (Fig. 1A, D). For model sharing, we can share all the maps, PDB files and 3D objects and the JSON file. As most of the components are from EMDB and PDB, the JSON is the only one needed to exchange the model that we created, which is very portable due to its small size.

### Case studies

#### *Chlamydomonas* cilium using density maps

To demonstrate CiliaBuilder2 capability, we build the *Chlamydomonas* cilia using *Chlamydomonas* maps of the doublet microtubule (EMD-2132) (Bui, Yagi et al. 2012) and central pair (EMD-31143) (Oda, Yanagisawa et al. 2014) (Fig. 2A). CiliaBuilder2 can also generate doublet microtubules in the same cilium with random offsets. In addition, a ciliary membrane with or without the cap covering the cilium can be also built (Fig. 2B).

**Fig. 2.**
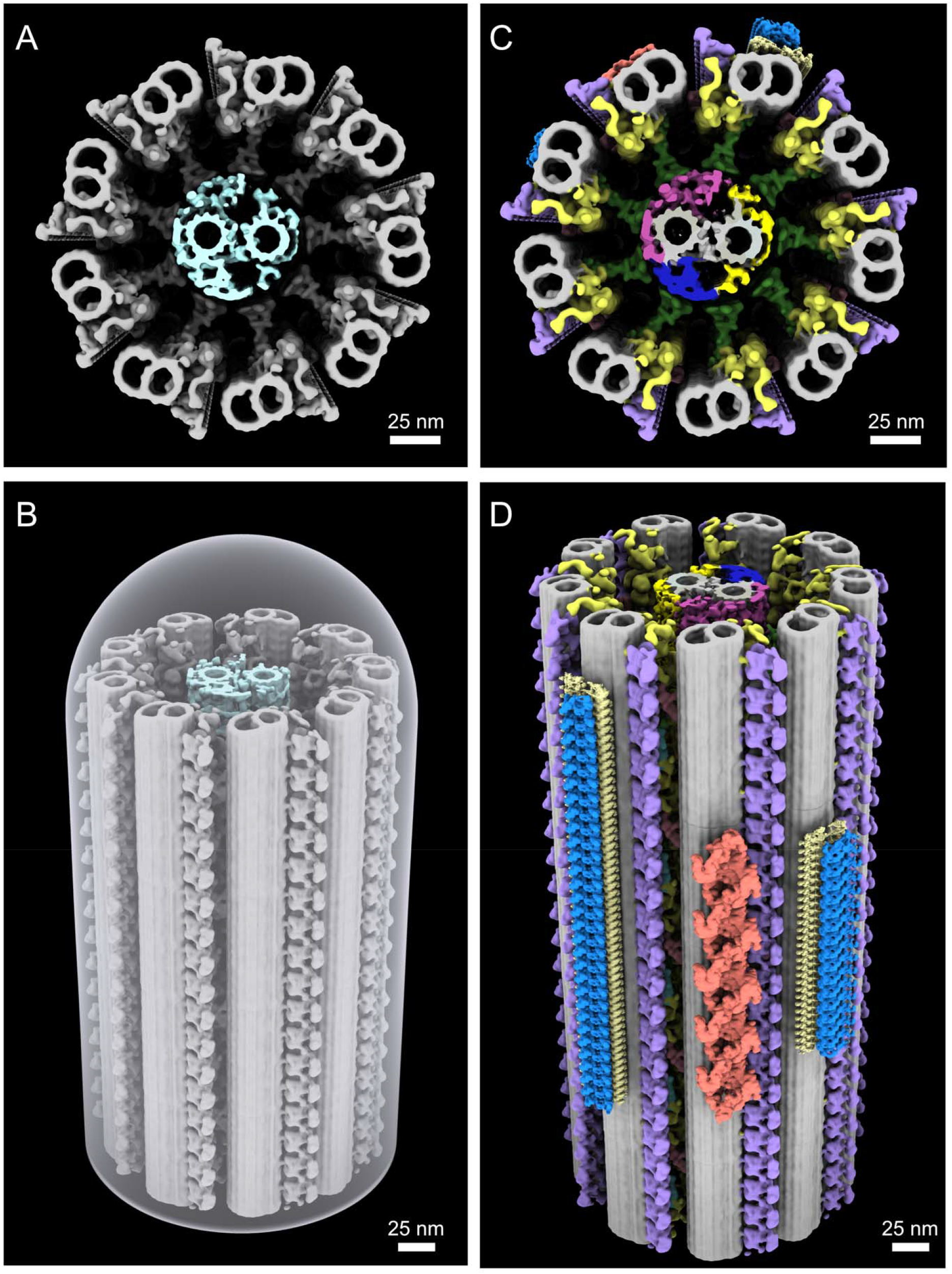
*Chlamydomonas* cilia built by CiliaBuilder2. (A) A simple *Chlamydomonas* cilium built from density map (Doublet microtubule EMD-2132 and central pair EMD-31143). (B) The same cilium as (A) with a ciliary membrane. (C) A *Chlamydomonas* cilium built using colored GLB files. (D) A *Chlamydomonas* cilium including anterograde and retrograde IFT built using colored GLB files (Anterograde IFT-B EMD-16014: blue; IFT-A EMD-15980: gold; Retrograde IFT EMD-19515: salmon). The IFT train is built by mimicking the figures from IFT publication (Lacey, Foster et al. 2023, Lacey, Graziadei et al. 2024).

#### *Chlamydomonas* cilium using colored GLB files

The previous example used maps, but each map can only be colored with a single color. While map coloring can be done using ChimeraX markers or segmentation before attachment, it is rather cumbersome to color and attach in the same session. CiliaBuilder2 can overcome this problem by using colored GLB files as attachment.

In this case, we used ChimeraX marker feature to color different components of the doublet microtubule (Outer dynein arm, inner dynein arm, radial spokes, nexin-dynein regulatory complex) and save the colored map as a single GLB file. For the central pair, we colored the protrusion differently from the C1 and C2 microtubules and saved the colored map as a single GLB file. Afterwards, the GLB files were attached, allowing a nice colored cilia model (Fig. 2C). As ChimeraX natively support reading of compressed GLB.GZ files natively, we compressed the GLB files before loading and significantly reduce the file size.

To show the flexibility of our program, we built the anterograde IFT using the compressed GLB models from IFT-A (EMD-15980), IFT-B (EMD-16014) (Lacey, Foster et al. 2023) and retrograde IFT (EMD-19515) (Lacey, Graziadei et al. 2024) by mimicking the figures from such studies (Fig. 2D).

### Sperm cilium

In this case, we built a fully complete PDB of a sperm cilium. We constructed a short cilium (2 × 96 nm repeat) using the doublet microtubule model from bovine sperm (PDB: 9fqr, 4527365 atoms) (Leung, Sun et al. 2025) and the central pair model (6 × 32 nm repeat) from mouse sperm (PDB: 9ijj, 1637265 atom) (Zhu, Lin et al. 2025).

This scene consists of over 50 million atoms, which astonishingly can be rendered reasonably quick on a MacBook Pro laptop with M1 Max 10-core CPU, 24-core GPU and 64 GB RAM. (Fig. 3).

**Fig. 3:**
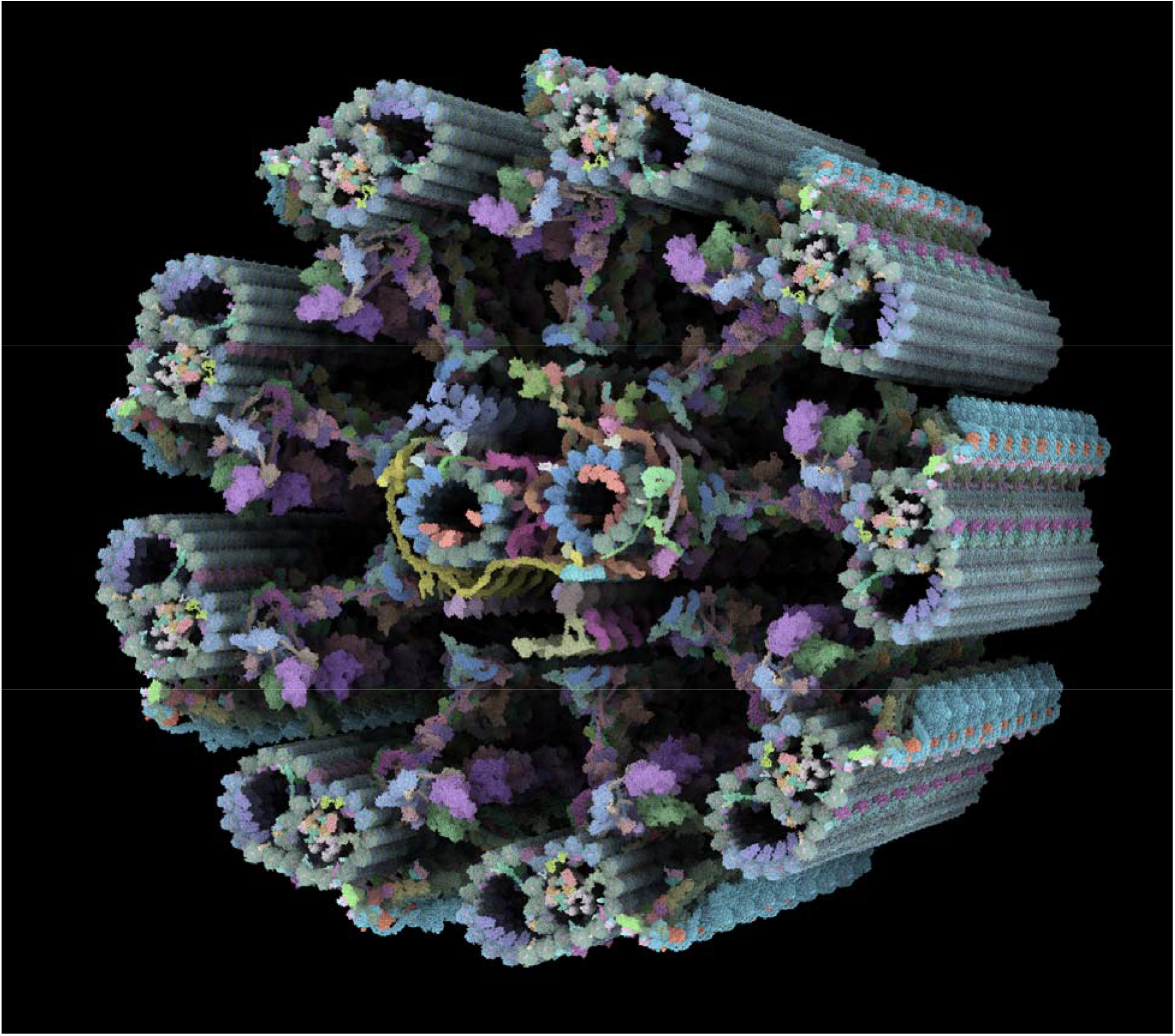
A hybrid model for a sperm cilium. The model is built from the bovine doublet microtubules (PDB: 9fqr) (Leung, Sun et al. 2025) and the mouse sperm central pair (PDB 9ijj) (Zhu, Lin et al. 2025).

### Centriole

We constructed a centriole model from *Tetrahymena*. The centriole consists of the proximal (EMD-42776), core (EMD-42777) and distal region (EMD-42778) (Ruehle, Li et al. 2024). We drew 3 microtubule coordinates consecutively, proximal (100 nm), core (300 nm) and distal region (100 nm) (Fig. 4).

**Fig. 4.**
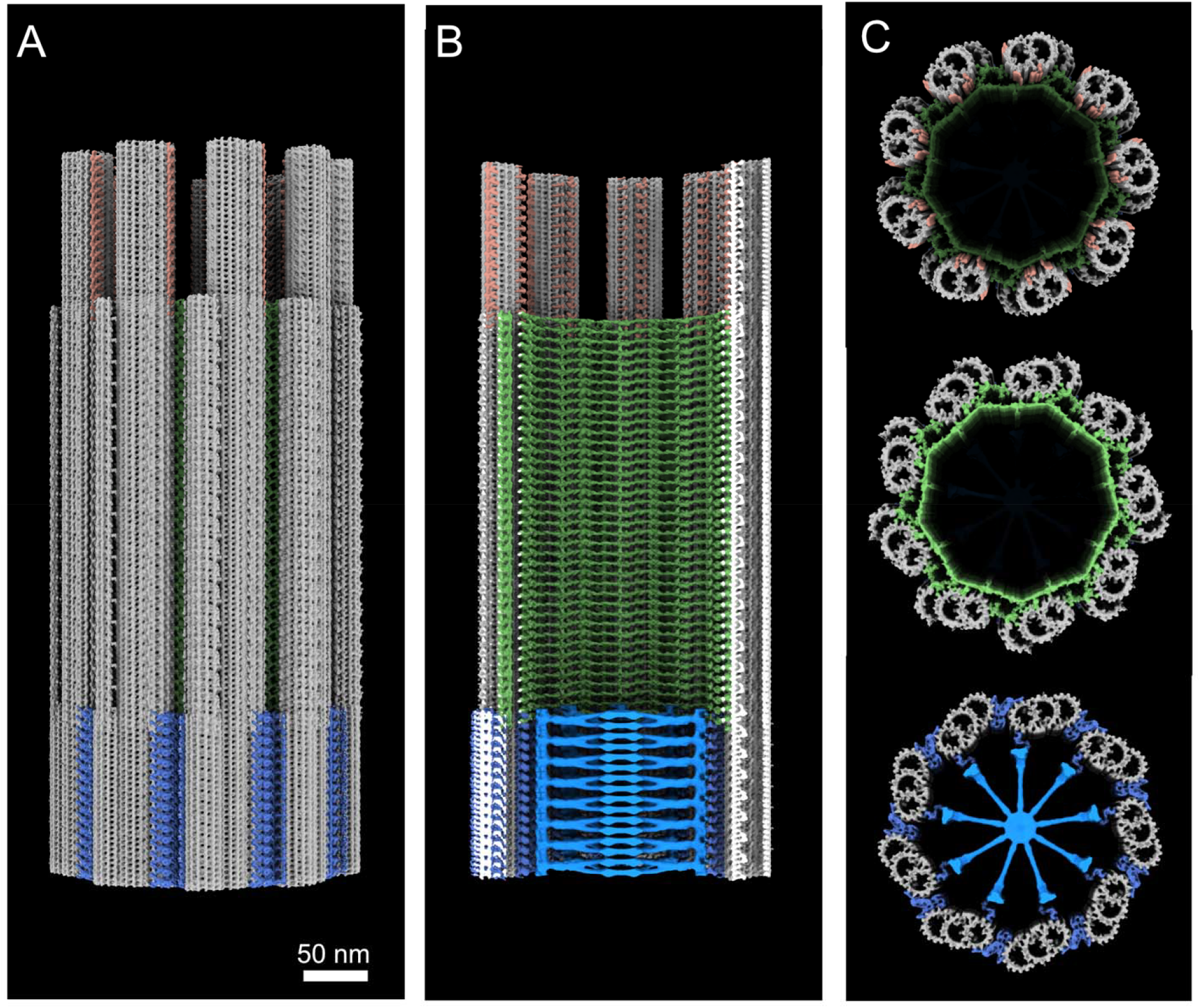
Tetrahymena centriole. (A) *Tetrahymena* centriole built from the proximal triplet microtubule (EMD-42776), core triplet microtubule (EMD-42777) and distal doublet microtubule (EMD-42778) (Ruehle, Li et al. 2024) and cartwheel (EMD-2329) (Guichard, Hachet et al. 2013). (B) A cut-in-half centriole view. (C) Cross sections of the distal (top), core (middle) and proximal (bottom) regions.

In addition, we generated the coordinate for the cartwheel for the proximal region and attach the cartwheel based on Trichonympha (EMD-2329) (Guichard, Hachet et al. 2013).

## Discussion

CiliaBuilder2 provides a convenient and flexible way to visualize cilia and centrioles. With the JSON file, we can also share either the session or the complete GLB so that the community can be benefited from models.

The centriole example in this study demonstrate that we can model ciliary longitudinal diversity with CiliaBuilder2. To model radial asymmetry, in which one or a few doublet microtubules are different from others (Bui, Sakakibara et al. 2009), CiliaBuilder2 provides the ability the delete selected coordinate points to create coordinates with less than nine filaments to attach the asymmetric doublet microtubules.

Apart from the fact that CiliaBuilder2 can work with not only maps but also GLB and PDB files, there are a few differences with ArtiaX. The main difference is on the original data taken in use: CiliaBuilder2 can generate parametric coordinate data and import the experimental data through STAR file, while ArtiaX only provides the interface for the original data file input with fixed orientations. This can cause several problems. The experimental data of the cilia can represent very short region or incomplete in the case of in situ cryo-Focus Ion Beam. In addition, cilia and centriole can be distorted due to the vitrification process. Furthermore, CiliaBuilder2 provides an easy way to tweak maps of different orientations to fit with the cilia and centriole geometry during the attachment process, allowing a flexible input map if the filament axis is parallel to Y or Z within the map.

While our example showed that CiliaBuilder2 can even work for a session containing over 50 million atoms, atomic models of the cilia present a computational challenge. Perhaps, exporting the CiliaBuilder2 scene in CellPack and Mesoscope format will allow efficient display as they use a better engine for display. This can be a potential improvement for the future version of CiliaBuilder2.

CiliaBuilder2 pioneered parametric modeling for straight ciliary architecture. Although CiliaBuilder2 currently lacks native support for generating curve cilia or geometries with variable diameters—such as those found in the ciliary transition zone—custom STAR files defining these geometric variations can be generated via scripting and imported directly into CiliaBuilder2 for map attachment. This approach can be extended to model other biological structures, including viruses and bacteria, thereby broadening the scope of available geometries. Furthermore, modeling complex cellular morphology and protein packing such as in membrane can be achieved by integrating CiliaBuilder2-like program with complementary tools such as CellPACK.

## Materials & Methods

### Example 1: *Chlamydomonas* cilium

Outer doublet microtubules were configured with a length of 3000 Å, a periodicity of 1000 Å, a radius of 960 Å, and nine doublet microtubules, maintaining a twist per layer of 0° and a Z offset of 0 Å. The central pair was defined with a length of 3000 Å, a periodicity of 333 Å, a Z offset of 0 Å, and constructed under Single line mode.

Density maps for doublet microtubule (EMD-2132) (Bui, Yagi et al. 2012) and central pair (EMD-31143) (Oda, Yanagisawa et al. 2014) were fetched directly from Electron Microscopy Data Bank inside ChimeraX. The EMD-2132 doublet map was fitted using a volume threshold of 1, with attachment rotations of 257° around the Z-axis and 0° around the Y-axis. Similarly, the EMD-31143 central pair map was placed using a volume threshold of 1, with attachment rotations of 257° around the Z-axis and 0° around the Y-axis.

The membrane was set to 4000 Å in length, 1350 Å in radius and Z offset of -700 Å with a thickness of 40 Å.

### Example 2: *Chlamydomonas* cilium using GLB

Density maps for EMD-2132 (Bui, Yagi et al. 2012) and EMD-31143 (Oda, Yanagisawa et al. 2014)) were individually segmented and colored using ChimeraX markers to distinguish outer dynein arms, inner dynein arms, radial spokes, the nexin-dynein regulatory complex, and central pair protrusions. The resulting scenes were exported as GLB files, setting the centering option to false to preserve the original map alignment.

The structural scaffold was constructed by defining nine outer doublet microtubules with a length of 4000 Å, a periodicity of 1000 Å, a radius of 960 Å, a twist per layer of 0°, and a Z offset of 0 Å. The central pair was modeled under Single line mode with a length of 3000 Å, a periodicity of 333 Å, and a Z offset of 0 Å. The corresponding GLB files were then integrated into the structural scaffold using the previously established attachment parameters using a fixed center of rotation (721.375, 721.375, 721.375) Å for the doublet microtubule and (614.688, 614.688, 614.688) Å for the central pair reading from the corresponding map. The use of the fixed rotation center allows attaching of GLB files with exact same building parameter as the maps.

For the IFT, retrograde map EMD-19515 (Lacey, Graziadei et al. 2024), anterograde IFT-B EMD-16014 (Lacey, Foster et al. 2023) and IFT-A EMD-15980 (Lacey, Foster et al. 2023) were fetched from Electron Microscopy Data Bank. EMD-19515 and EMD-15980 were aligned their long axis along z manually and saved as corresponding GLB files.

We modelled two anterograde IFT trains. For the first anterograde IFT train positioned on microtubule doublet (filament) 3, IFT-B (EMD-16014) was modeled at a radial distance of 1250 Å from the center with an anterograde angle of -6°, an orientation angle of 0°, a Z offset of 1500 Å, a periodicity of 65 Å, 25 repeat units, and a 180° Y-axis attachment rotation with no fixed rotation center. On the same doublet, IFT-A (EMD-15980) was modeled at a radial distance of 1380 Å from the center with an anterograde angle of 3°, an orientation angle of 0°, a Z offset of 1500 Å, a periodicity of 115 Å, 14 repeat units, and an attachment setting of 0° for both Z and Y rotations with no fixed rotation center. A second anterograde IFT train was placed on filament 1 using equivalent spatial parameters, with an adjusted Z offset of 1000 Å and repeat counts of 50 for IFT-B and 28 for IFT-A.

The retrograde IFT train (GLB file) was positioned on filament 2 at a radial distance of 1300 Å from the center, featuring a retrograde angle of 0°, a Z offset of 1700 Å, a periodicity of 900 Å, 2 repeat units, and attachment rotations of 190° around the Z-axis and 90° around the Y-axis with no fixed rotation center.

### Example 3: Sperm cilium

Outer doublet microtubules were configured using a length of 1900 Å, a periodicity of 1000 Å, a radius of 800 Å, and nine doublets, with a twist per layer of 0° and a Z offset of 0 Å to generate 2 times 96 nm repeats. The central pair was specified in “Single line” mode with a length of 1900 Å, a periodicity of 333 Å, and a Z offset of -333 Å.

Atomic structures for the bovine doublet microtubule (PDB: 9FQR) (Leung, Sun et al. 2025) and mouse central pair (PDB: 9IJJ) (Zhu, Lin et al. 2025) were retrieved directly from the Protein Data Bank within UCSF ChimeraX. Both models were attached to the scaffold with 0° rotation around both the Z- and Y-axes.

### Example 4: *Tetrahymena* Centriole

For the *Tetrahymena* centriole, three consecutive segments were constructed with lengths of 1000 Å, 3000 Å, and 1000 Å. All three segments shared identical geometric parameters: a periodicity of 165 Å, a radius of 960 Å, nine triplet microtubules, a 0° twist per layer, and a Z offset of 0 Å. The cartwheel line was defined along the central pair axis with a length of 835 Å, a periodicity of 165 Å, and a Z offset of -82.5 Å.

Cryo-EM maps representing the proximal (EMD-42776), core (EMD-42777), and distal (EMD-42778) (Ruehle, Li et al. 2024) regions were retrieved from the Electron Microscopy Data Bank (EMDB). The proximal map was first aligned so that the microtubule axis ran parallel to the Z-axis, and the oriented volume was saved. Subsequently, the core and distal maps were aligned, resampled relative to the oriented proximal map, and exported. Using CiliaBuilder2, the center of rotation for the three maps are calculated to be (564.300, 564.300, 564.300) Å.

Non-tubulin density from three maps were colored at a threshold of 0.122 and the resulting color surfaces were saved to GLB individually with the false centering.

All three GLB files were attached across their respective consecutive regions using a -8° Z-axis rotation and a 0° Y-axis rotation and a fixed center of rotation of (564.300, 564.300, 564.300) Å. Finally, the cartwheel density map (EMD-2329) (Guichard, Hachet et al. 2013) was fetched, converted to GLB at level of -3.07 and attached onto the central line with 20° rotation along Z-axis.

## Contributions

T.L. coding, K.H.B. idea, supervision, S.C. & K.H.B testing. T.L., S.C. and K.H.B writing.

## Acknowledgement

K.H.B. is supported by grants from the Centre de recherche en biologie structurale Bluesky Seed Funding, Canadian Institutes of Health Research (PJT-190195) and the Natural Sciences and Engineering Research Council of Canada (RGPIN-2022-04774).

## Disclosure of AI use

The authors declare the use of generative AI (GAI) in the research and writing process. The following tasks were delegated to GAI tools under full human supervision: code generation (Claude Opus 5.5), proofreading and editing (Gemini 3.0).

## Data Availability

CiliaBuilder2 can be installed directly from the Toolshed within UCSF ChimeraX. https://cxtoolshed.rbvi.ucsf.edu/apps/chimeraxciliabuilder2

The code for CiliaBuilder2, JSON files and associated files and guideline for CiliaBuilder2 are available on: https://github.com/builab/CiliaBuilder2

